# BRD4 regulates Aurora B kinase activity

**DOI:** 10.1101/2025.08.06.668960

**Authors:** Ballachanda N. Devaiah, Dan Cheng, Amit K. Singh, Dinah S. Singer

## Abstract

BRD4, a pleiotropic regulator of chromatin structure and transcription, plays critical roles in cancer and immune responses. Unlike other transcriptional regulators, BRD4 largely remains bound to chromosomes during early mitosis. Here we report that BRD4 also regulates mitosis through its direct interaction with and phosphorylation of Aurora B kinase, an essential regulator of mitosis. BRD4 binding to Aurora B inhibits its kinase activity, preventing autophosphorylation and phosphorylation of the key mitotic targets histone H3 and MCAK, the mitotic centromere associated kinesin. This inhibition is relieved during metaphase when JNK is activated and phosphorylates BRD4, triggering its transient release from chromatin. Importantly, Aurora B activity during mitosis inversely correlates with BRD4 binding and directly correlates with JNK activation and BRD4 release. Our findings thus reveal a regulatory mechanism whereby Aurora B activity is directly controlled by BRD4, which in turn is regulated by JNK.

**Significance Statement:** BRD4 has been extensively characterized for its role in regulating chromatin structure and transcription. But its function during mitosis has remained unclear. This study reveals a novel mechanism by which BRD4 directly regulates mitotic progression through its interaction with and inhibition of Aurora B kinase, a central player in chromosome segregation. The timely release of BRD4 from chromatin via JNK-mediated phosphorylation enables Aurora B activation at a critical stage of mitosis. These findings uncover a previously unrecognized BRD4–Aurora B–JNK signaling axis that integrates chromatin dynamics with mitotic control, offering new insights into cell cycle regulation and potential therapeutic targets in cancer.

## Introduction

The Aurora B kinase is a mitosis regulator that plays a critical role in chromosomal stability, segregation and cytokinesis through its phosphorylation of numerous key players during mitosis (1). Aurora B gene expression is strongly induced in the G2 phase of cell cycle with high levels of Aurora B protein observed during early mitosis/prophase through the end of mitosis (1). Aurora B kinase is activated by autophosphorylation of its threonine 232 (T232) site (2). However, despite its presence in all stages of mitosis, Aurora B kinase exhibits peak activity during the anaphase and telophase stages that follow metaphase (3), suggesting that its kinase activity is suppressed prior to anaphase.

BRD4 is a BET family bromodomain protein with pleiotropic functions that include chromatin and transcriptional regulation, alternative RNA splicing, DNA repair and cell cycle regulation (4–6). BRD4 has both intrinsic histone acetyltransferase (HAT) and kinase activities through which it regulates chromatin architecture and transcription respectively (7, 8). We recently reported that JNK-mediated phosphorylation releases BRD4 from chromatin and its role as a chromatin modifying HAT and activates its transcriptional kinase (9). JNK activation and BRD4 transcriptional activity are significantly enhanced during cellular stress, immune/inflammatory responses and epithelial to mesenchymal transition (EMT) during cancer progression. Notably, JNK activation has also been reported to surge in the metaphase stage during mitosis (10). Unlike most transcriptional and chromatin regulators, BRD4 remains associated with the chromatin through most of mitosis but is briefly released after metaphase (11), concurrent with JNK activation. While the presence of BRD4 on mitotic chromatin has been proposed to serve as a passive bookmark for post-mitotic transcriptional activity (12, 13), the precise reason for its strong association and brief release is yet to be fully elucidated.

In this study, we hypothesized that BRD4 remains associated with mitotic chromatin in part to suppress Aurora B activity. Indeed, we report that BRD4 directly interacts with Aurora B kinase on chromatin. This interaction results in the inhibition of Aurora B kinase activity and the subsequent phosphorylation of its substrates, histone H3 and the mitotic centromeric associated kinesin (MCAK), critical for mitotic progression. Strikingly, phosphorylation of BRD4 by activated JNK leads to BRD4’s release from chromatin and abrogates its inhibition of Aurora B kinase activity. Accordingly, BRD4 and Aurora B co-localize on chromatin during the early stages of mitosis when Aurora B activity is attenuated but are no longer co-localized after metaphase when JNK is activated and BRD4 has been released from chromatin. Based on these results we propose a model whereby Aurora B activity is directly regulated by BRD4 and indirectly by JNK activation during mitosis.

## Results

### BRD4 and Aurora B directly interact on chromatin

Since both BRD4 and Aurora B are associated with mitotic chromatin (1, 11), we first asked if the two proteins interact. Indeed, Aurora B robustly co-immunoprecipitated with BRD4 from HeLa nuclear extracts, indicating that BRD4 and Aurora B exist in a complex in the nucleus (Fig. 1A). This interaction occurs in vivo, as shown by a proximity ligation assay (PLA) in HCT116 cells which showed that BRD4 and Aurora B exist in close proximity (Fig. 1B). In an in vitro pull-down assay, recombinant purified His-BRD4 bound to Ni-NTA beads efficiently pulled down increasing amounts of recombinant GST-Aurora B, demonstrating that the interaction between BRD4 and Aurora B is direct (Fig. 1C). BRD4’s kinase domain, as well as its two bromodomains are located in its N-terminal region; its acetyl CoA binding site and HAT catalytic site are in its C-terminal end (14). Interestingly, the deletion of BRD4’s bromodomains, or either its N- or C-terminal segments did not eliminate Aurora B binding, suggesting that Aurora B binds to multiple alternative sites on BRD4 (Fig. 1D). While small quantities of BRD4 and Aurora B are chromatin-free, the BRD4-Aurora B interaction is detectable only on the chromatin-bound cell protein fraction. Thus, BRD4 and Aurora B co-immunoprecipitate from an extract derived from the chromatin-bound fraction but not from a chromatin-free fraction of HCT116 cell extracts (Fig. 1E). Together, these results demonstrate that BRD4 and Aurora B interact directly on chromatin.

**Fig. 1.**
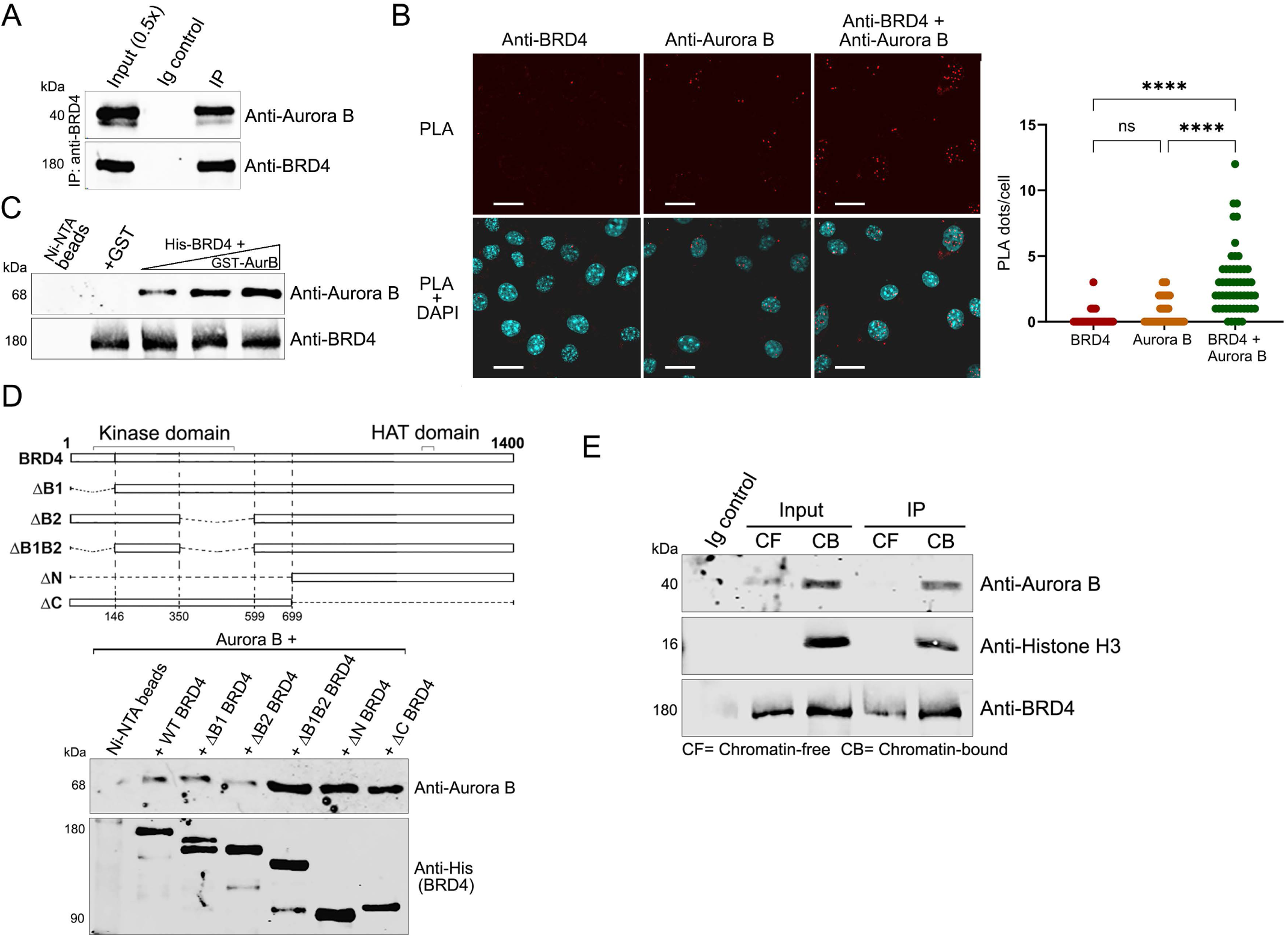
BRD4 and Aurora B interact on the chromatin. **A.** BRD4 associates with Aurora B. BRD4 was immunoprecipitated from HeLa nuclear extract using anti-BRD4 and immunoblotted with anti-Aurora B. **B.** BRD4 co-localizes with Aurora B in the nucleus. (Left) Proximity ligation assays (PLA) with anti-BRD4 and anti-Aurora B on fixed HCT116 cells. PLA-red; DAPI nuclei staining-blue. Scale bars; 25 µM. (Right) Quantification of PLA shown on left comparing the BRD4-Aurora B proximity with BRD4 and Aurora B alone antibody controls. **C.** BRD4 binds Aurora B directly. GST-Aurora B (0.15, 0.2 and 0.25µg) was pulled down with 0.5µg His-BRD4 immobilized on Ni-NTA beads. Beads with GST protein or without prey protein are controls. **D.** Aurora B interacts with multiple regions on BRD4. Top: Map of BRD4 and deletion mutants. Bottom: anti-Aurora B immunoblot of 0.15 µg Aurora B recovered by pull-down with 0.5 µg wild type His-BRD4-Flag or equimolar amounts of His-BRD4-Flag mutants on Flag beads. Anti-His immunoblot shows BRD4 retained on Flag-beads. **E.** Aurora B interacts with chromatin-bound BRD4. Immunoblots show Aurora B co-immunoprecipitating with BRD4 immunoprecipitated from chromatin-free and chromatin-bound protein fractions of HCT116 cells with anti-BRD4 antibody.

### BRD4 suppresses Aurora B phosphorylation of Histone H3 and MCAK

Based on our finding that BRD4 and Aurora B kinases interact, we next tested whether that interaction affects Aurora B kinase activity. The substrates of Aurora B kinase include histone H3, whose phosphorylation is a hallmark of mitosis, and the mitotic centromere associated kinesin (MCAK), the phosphorylation of which is critical for segregation of chromosomes during mitosis and safeguarding chromosome stability. We first examined the effect of BRD4 on phosphorylation of histone H3 by Aurora B. There are three main variants of histone H3: H3.1, H3.2 and H3.3. While Histone H3.1 and H3.2 are canonical H3 variants primarily associated with transcriptionally silent heterochromatin regions, H3.3 is the replacement histone associated with transcriptionally active gene loci on euchromatin. H3.3 is also the most abundant H3 variant found on chromatin. In an in vitro kinase assay, Aurora B differentially phosphorylated the three histone H3 isoforms H3.1, H3.2 and H3.3 isoforms in hyper- and hypo-phosphorylation patterns (Fig. 2A). BRD4 did not phosphorylate any of the H3 isoforms. Importantly, the presence of BRD4 in the kinase assay significantly affected Aurora B phosphorylation of the three H3 isoforms, although to different degrees. BRD4 inhibited Aurora B hyper-phosphorylation of H3.1 and H3.3, but had no effect on Aurora B phosphorylation of histone H3.2 (Fig.2A). Thus, BRD4 selectively modulates Aurora B phosphorylation of the histone H3 variants.

**Fig. 2.**
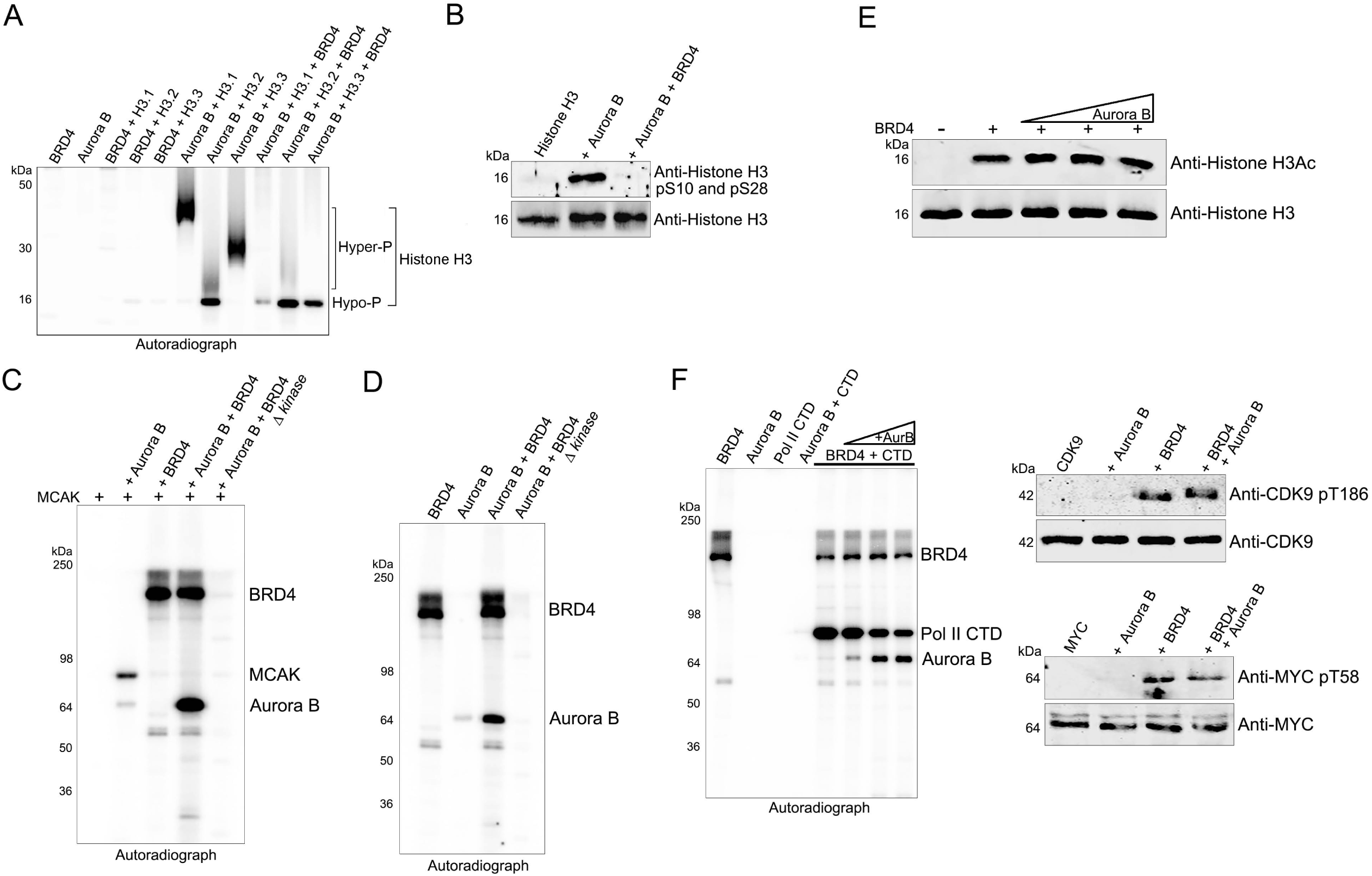
BRD4 phosphorylates and inhibits Aurora B. **A.** BRD4 inhibits Aurora B phosphorylation of specific histone H3 isoforms. Autoradiograph of kinase assays with equimolar Aurora B, 0.3µg BRD4 and histone H3.1, H3.2 and H3.3 as substrates. **B.** BRD4 inhibits Aurora B phosphorylation of histone H3 Ser10 and Ser28. Immunoblots of kinase assays done with equimolar Aurora B, 0.3µg BRD4 and histone H3.1, using a combination of antibodies targeting histone H3 pS10 and pS28. **C.** BRD4 inhibits Aurora B phosphorylation of MCAK. Autoradiograph of kinase assays with equimolar Aurora B, BRD4 and BRD4 Δ kinase with MCAK as substrate. **D.** BRD4 phosphorylates Aurora B. Autoradiograph of kinase assays with equimolar Aurora B and 0.3µg BRD4 WT or BRD4 Δ kinase. **E.** Aurora B does not affect BRD4 HAT activity. Immunoblots of a HAT assay in the presence of ATP with 0.3µg BRD4 and equimolar histone H3 without or with increasing amounts (0.1, 0.2, 0.3 and 0.4µg) of Aurora B. **F.** Aurora B does not directly affect BRD4 kinase activity. Left: Autoradiograph of kinase assays with BRD4 and equimolar or increasing amounts of Aurora B with Pol II CTD as substrate. Right: Immunoblots of kinase assays done with equimolar Aurora B and BRD4 with pTEFb/CDK9 (top panel) or MYC (bottom panel) as substrates.

Chromosome condensation and segregation during mitosis depend on phosphorylation of histone H3 specifically at Ser10 and Ser28 by Aurora B (15). In an in vitro kinase assay, Aurora B phosphorylation of histone H3.3 Ser10 and Ser 28 was completely abrogated in the presence of BRD4, as assessed by immunoblotting with an antibody specific for H3pS10 and pS28 (Fig. 2B). Mitosis further depends on another critical Aurora B kinase substrate, MCAK, which contributes to mitotic spindle formation. Accordingly, in vitro, Aurora B, but not BRD4, efficiently phosphorylated MCAK (Fig. 2C). Importantly, BRD4 inhibited phosphorylation of MCAK by Aurora B. (Fig. 2C, lanes 2,4). Taken together, these findings indicate that BRD4 contributes to the regulation of mitosis through its direct interaction with and inhibition of Aurora B kinase activity.

### BRD4 kinase activity is not required for the suppression of Aurora B kinase activity

While BRD4 suppressed the kinase activity of Aurora B, it also phosphorylated Aurora B (Fig. 2C, lane 4). This led to question of whether the suppression of Aurora B kinase activity by BRD4 depended on BRD4’s kinase activity. Surprisingly, a kinase-dead mutant of BRD4, BRD4 Δ kinase (16), still suppressed Aurora B phosphorylation of MCAK (Fig. 2C, lane 5). Further, BRD4 kinase activity is necessary to phosphorylate Aurora B, in the presence or absence of MCAK. As shown in Fig. 2D, BRD4 but not the kinase-dead mutant, phosphorylates Aurora B. The role of this phosphorylation in regulating Aurora B remains to be determined. Aurora B does not phosphorylate BRD4, as evidenced by the lack of phosphorylation of the BRD4 Δ kinase mutant (Fig. 2C and 2D).

We further asked whether Aurora B regulates BRD4’s HAT and kinase activities. In an in vitro HAT assay in the presence of ATP, BRD4 acetylation of histone H3 remained unaffected by increasing amounts of Aurora B, indicating that Aurora B does not affect BRD4 HAT activity (Fig. 2E). BRD4 phosphorylation of the RNA Pol II carboxyterminal domain (CTD) was only minimally affected by Aurora B at equimolar concentrations, with a slight reduction in CTD phosphorylation observed at high concentrations of Aurora B, presumably through competitive inhibition by Aurora B (Fig. 2F; Left panel). Similarly, BRD4 phosphorylation of CDK9 Thr186 and of MYC Thr58 was unaffected by Aurora B (Fig. 2F; Right panels). These results thus demonstrate that Aurora B does not markedly affect BRD4 enzymatic activities.

### JNK activation promotes Aurora B activity in vivo through BRD4 release

We next investigated how BRD4 suppression of Aurora B kinase activity could be relieved to allow mitotic progression. One possible explanation was that BRD4 phosphorylation of Aurora B disrupted their interaction. This possibility was excluded by the finding that following phosphorylation of Aurora B by BRD4 *in vitro*, BRD4 and Aurora B still efficiently coimmunoprecipitated (Fig 3A).

**Fig. 3.**
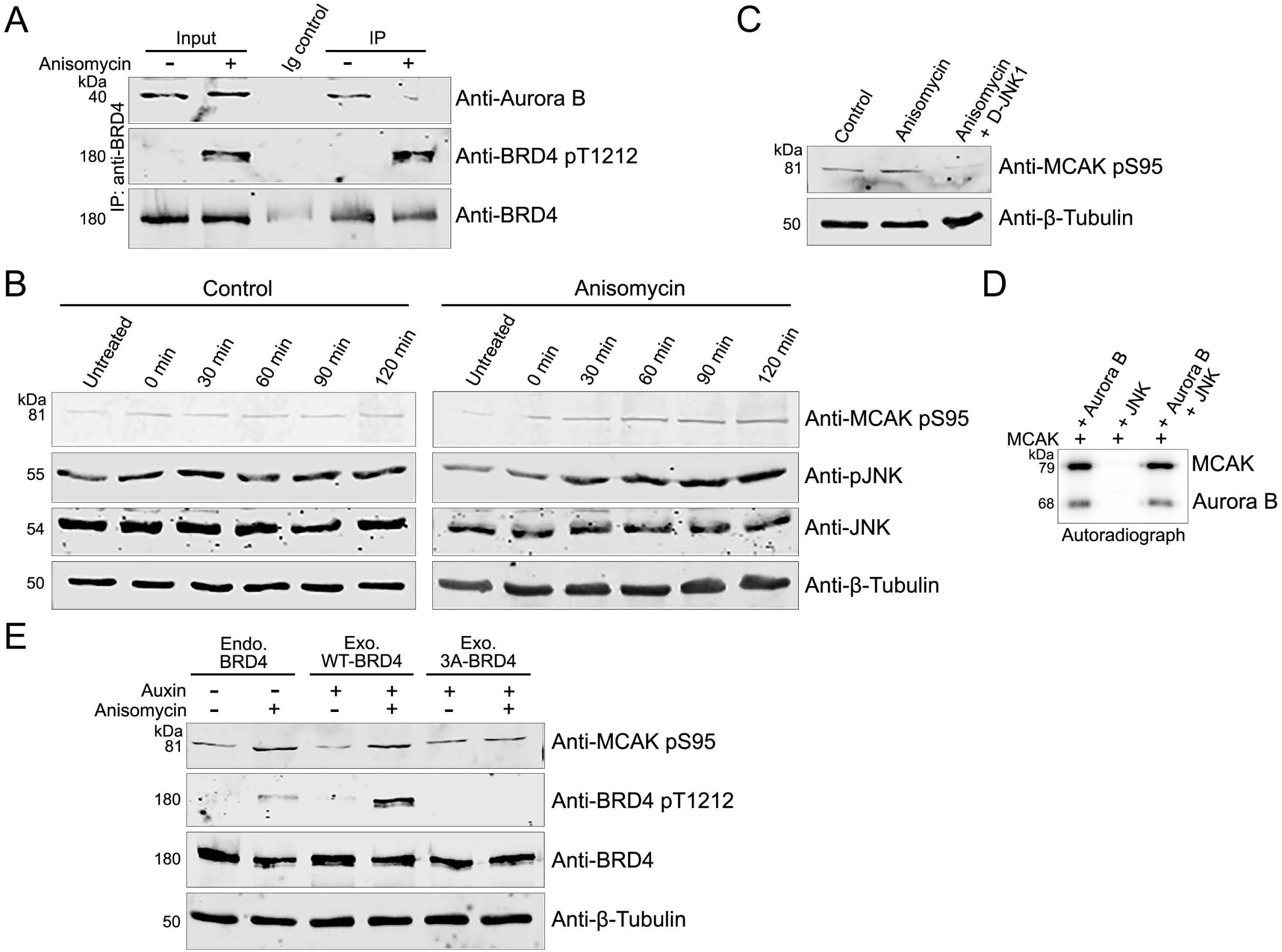
JNK modulates BRD4 regulation of Aurora B. **A.** Aurora B interaction with BRD4 is unaffected by its phosphorylation. Co-IP of Aurora B with Flag-BRD4 on Flag beads following a kinase assay with or without ATP. **B.** JNK activation and phosphorylation of BRD4 *in vivo* abrogates BRD4-Aurora B interaction. Immunoblots showing Aurora B co-immunoprecipitated by anti-BRD4 with total and T1212 phosphorylated BRD4 from HCT116 cells treated with or without JNK activator anisomycin. **C.** JNK activation increases MCAK Ser95 phosphorylation. Immunoblots showing MCAK pSer95 and pJNK levels in HCT 116 cells with or without JNK activation by anisomycin for increasing durations of time. Data is representative of two independent experiments. **D.** JNK inhibition decreases MCAK Ser95 phosphorylation. Immunoblots showing MCAK pSer95 levels in HCT116 cells with or without anisomycin and JNK inhibitor D-JNK1. **E.** JNK does not directly phosphorylate MCAK or affect Aurora B activity. Autoradiograph of kinase assays with equimolar Aurora B and active JNK with MCAK as substrate. **F.** MCAK phosphorylation is dependent on JNK phosphorylation of BRD4. Immunoblots showing MCAK pSer95 levels in DLD-BRD4-IAA7 cells transfected with or without WT or 3A-BRD4 and treated with or without auxin and anisomycin.

We recently reported that JNK phosphorylates BRD4, releasing it from chromatin (9). Since JNK is activated during mitosis, we considered the possibility that BRD4 phosphorylation by JNK abrogates its interaction with Aurora B. As we have shown previously, activation of JNK by anisomycin treatment of HCT116 cells increased JNK phosphorylation of BRD4 at Thr1212 (9). This increased phosphorylation of BRD4 correlated with a concomitant decrease in the co-immunoprecipitation of Aurora B with BRD4 (Fig. 3B). We predicted that the JNK-mediated phosphorylation of BRD4, which led to a reduced interaction between BRD4 and Aurora B, would result in increased Aurora B kinase activity in vivo. To test this, JNK was activated with anisomycin for increasing times; levels of MCAK phosphorylation at Ser95, an Aurora B specific target site, were assessed (Fig. 3C). Cells pre-treated with the JNK inhibitor SP600125 and then treated with anisomycin for similar times served as controls. Strikingly, immunoblots of whole cells extracts (WCE) with an anti-MCAK pSer95 antibody showed increasing Aurora B phosphorylation of MCAK concomitant with increasing JNK activation as assessed by pJNK levels (Fig. 3C). In contrast, MCAK pSer95 and pJNK levels remained unchanged in the control cells. As SP600125 has been recently reported to inhibit kinases other than JNK, we confirmed these results using D-JNK1, a highly specific cell permeable peptide JNK inhibitor (Fig. 3D). Importantly, JNK does not directly phosphorylate MCAK, nor does it affect Aurora B phosphorylation of MCAK in in vitro kinase assays (Fig. 3E). Therefore, JNK activation of Aurora B kinase is indirect but can be blocked by inhibiting JNK. These results thus demonstrate that JNK activation indirectly leads to Aurora B kinase activation and a consequent increase in MCAK phosphorylation.

To determine whether the increase in Aurora B activity upon JNK activation is a direct consequence of JNK phosphorylation of BRD4, we asked whether mutation of JNK’s phosphorylation site on BRD4 would abrogate JNK-mediated activation of Aurora B kinase. To this end, we used a DLD1-BRD4-IAA7 cell line where endogenous BRD4 is tagged with an auxin inducible degron and can be rapidly degraded by auxin treatment (17). Cells were transfected with or without a BRD4 mutant lacking JNK phosphorylation sites (3A-BRD4) or exogenous WT BRD4 and cultured in the presence or absence of anisomycin and auxin (Fig. 3F). Immunoblots of WCE from these cells showed an increase in MCAK pSer95 levels in the cells expressing either endogenous or exogenous WT BRD4, following JNK activation. In contrast, MCAK pSer95 levels remained unchanged in the cells expressing the exogenous mutant 3A-BRD4 following JNK activation (Fig. 3F). Further, the increase in MCAK phosphorylation correlated with JNK phosphorylation of WT BRD4 at Thr1212, which leads to a decreased interaction between BRD4 and Aurora B (Fig. 3B). Thus, the increase in MCAK phosphorylation by Aurora B upon JNK activation occurs through JNK phosphorylation-mediated release of BRD4 from Aurora B.

### BRD4-Aurora B colocalization on chromatin correlates with mitotic progression

Our results demonstrate that Aurora B kinase is suppressed by its interaction with BRD4; the suppression is alleviated when BRD4 is phosphorylated by JNK, which dissociates it from Aurora B (Figs. 1-3). Both Aurora B and JNK kinase are actively regulated during mitosis. Aurora B levels are maximum during prophase and metaphase, while JNK kinase activity peaks at metaphase (10), Therefore, we investigated whether the co-localization of BRD4 and Aurora B correlated with the stages of mitosis at which JNK and Aurora B kinases are activated. Immunofluorescence assays using specific antibodies were done to localize BRD4 and Aurora B at different stages of the cell cycle. As shown in Fig. 4A, while both BRD4 and Aurora B are expressed throughout mitosis, they are only colocalized during prophase when Aurora B kinase is suppressed (Fig. 4A; Upper panels). Strikingly, at metaphase when JNK activity peaks, BRD4 no longer co-localizes with Aurora B; rather it appears diffusely across the nucleoplasm while Aurora B remains concentrated on condensed metaphase chromatin (Fig. 4A; Middle panels). Following metaphase, during anaphase and telophase, Aurora B kinase becomes activated. At the same time, BRD4 re-localizes to chromatin but it is not colocalized with Aurora B (Fig. 4A; Lower panels). Therefore, co-localization of BRD4 and Aurora B correlates with both JNK and Aurora B kinase activities during the different stages of mitosis. These results extend our finding that BRD4 interacts with and directly suppresses Aurora B kinase activity; JNK activation relieves this suppression by phosphorylating BRD4, abrogating its interaction with Aurora B.

**Fig. 4.**
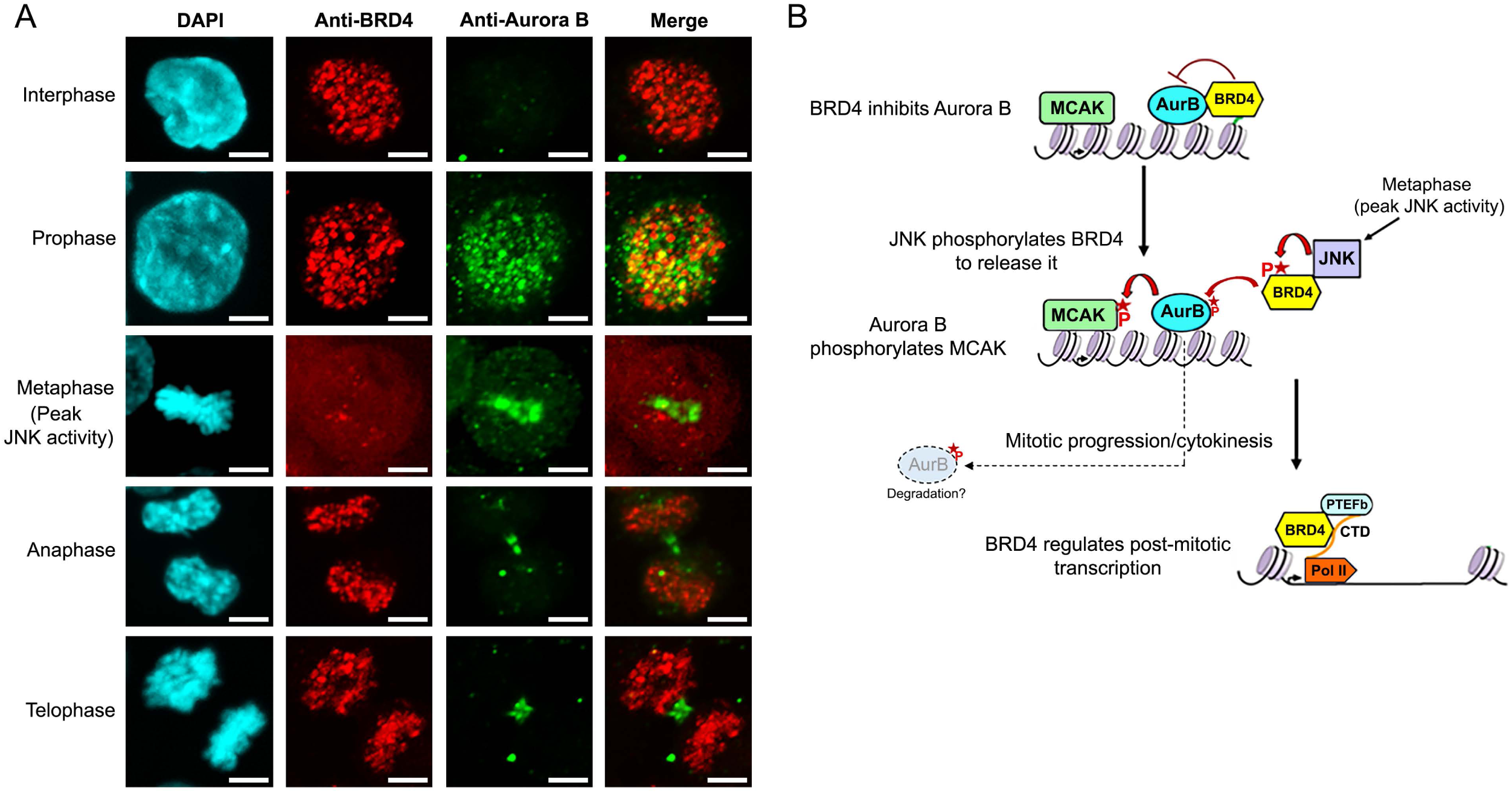
Mitotic progression is correlated with BRD4-Aurora B colocalization. **A.** BRD4-Aurora B co-localize in early mitosis but separate following metaphase. Immunofluorescence images showing BRD4 and Aurora B localization in HCT116 cells during different stages of the cell cycle. Scale bars; 5 µM. **B.** Model of BRD4 regulation of Aurora B during mitosis. BRD4 suppresses Aurora B kinase activity during early mitosis. During metaphase, elevated JNK activity causes the release of BRD4 from chromatin through its phosphorylation. The loss of BRD4 from chromatin and its separation from Aurora B leads to the reactivation of Aurora B kinase activity in anaphase and telophase. Aurora B is consequently able to phosphorylate MCAK and its other substrates allowing mitotic progression and cytokinesis. BRD4 released from chromatin by JNK phosphorylation activates post-mitotic transcription. BRD4 phosphorylation of Aurora B could possibly lead to its degradation.

## Discussion

Aurora B is a serine/threonine kinase that plays crucial roles during mitosis and cytokinesis, ensuring accurate chromosome segregation. Although Aurora B is expressed throughout the cell cycle, its kinase activity is primarily active during mitosis. The mechanisms regulating its kinase activity have not been extensively characterized. Our current studies demonstrate that Aurora B kinase activity is regulated by the bromodomain protein, BRD4, which interacts with Aurora B to inhibit its activity. This inhibition of Aurora B kinase activity is relieved during metaphase following JNK-mediated phosphorylation of BRD4 which results in the transient release of BRD4 from chromatin and the disruption of its interaction with Aurora B. During anaphase, Aurora B binds to the mitotic spindle whereas BRD4 reassociates with chromatin and is no longer co-localized with Aurora B.

The presence of BRD4 on mitotic chromatin, unlike other transcriptional regulators, has thus far been attributed to its possible role as a passive mitotic bookmark for transcription (12, 13), However, more recent reports have demonstrated its rapid transient release from chromatin at the end of mitosis (11), and confirmed that the removal of BRD4 does not impair post-mitotic activation of transcription (18). The current results reveal that BRD4 plays an active role during mitosis by modulating mitotic progress through its direct regulation of the Aurora B kinase. BRD4 has been previously reported to regulate the levels of Aurora B through transcriptional (19) and chromatin (20) regulation. The current findings showing post-transcriptional regulation of Aurora B by BRD4 identify an additional level of control by BRD4, extending BRD4’s known pleiotropic functions (14).

While small quantities of both BRD4 and Aurora B exist in chromatin-free states, PLA and biochemical fractionation indicate that detectable interaction occurs when both proteins are chromatin-bound (Fig. 1), suggesting that in cells BRD4 regulation of Aurora B is restricted to the association of both molecules with chromatin. We speculate this spatio-temporal BRD4 regulation of Aurora B could have important implications, as Aurora B also regulates key proteins such as p53, and cell processes like cytokinesis, in its chromatin-free state (21, 22).

In vivo, Aurora B is a component of the chromosomal passenger complex (CPC) - consisting of Aurora B, INCEP, Survivin and Borealin – that regulates the entire mitotic process (23). Activation of Aurora B kinase activity within the CPC has been shown to depend on its interaction with INCEP. The inhibition of Aurora B kinase by BRD4 in an *in vitro* assay with purified recombinant proteins demonstrates that the mechanism of inhibition is direct and independent of INCEP or the other CPC components. Although BRD4 phosphorylates Aurora B kinase, that is not the mechanism by which BRD4 inhibits Aurora B kinase activity: a kinase-dead BRD4 mutant is as efficient as the wild type BRD4 at inhibiting Aurora B kinase activity. Interestingly, Aurora B interacts with BRD4 in two separate domains located in the N-terminal and C-terminal halves of the BRD4 protein. Since BRD4 primarily exists as an antiparallel dimer (16) and Aurora B is known to form antiparallel dimers in solution (24), we speculate that dimeric Aurora B interacts with dimeric BRD4 to sequester Aurora B’s active site. Surprisingly, however, Aurora B does not affect either the kinase or HAT activity of BRD4, indicating that the interaction sites are distinct from BRD4’s enzymatic active sites.

The role of BRD4 phosphorylation of Aurora B is not known. BRD4-mediated phosphorylation does not affect Aurora B kinase activity, nor does it cause dissociation of the BRD4/Aurora B interaction. One possibility is that BRD4 phosphorylation of Aurora B enhances its interaction with INCEP, which is required for full kinase activity, or for its interactions with various substrates during mitosis (3). Alternatively, BRD4 phosphorylation could also serve as a mark for phosphorylation dependent FBXW7 mediated proteolytic degradation of Aurora B at the end of mitosis (25).

Aurora B kinase has numerous substrates whose phosphorylation is critical for mitotic progress, including Histone H3 and MCAK (15, 26); BRD4 inhibits Aurora B phosphorylation of MCAK and of two of three histone H3 variants. It completely blocks Aurora B phosphorylation of H3.3 S10 and S28 sites. In addition, BRD4 appears to specifically inhibit the hyper-phosphorylation of H3.1 and H3.3 by Aurora B without affecting the hypo-phosphorylation of H3.2. These differential effects of BRD4 on Aurora B kinase are surprising considering the highly conserved sequences of H3 and indicate that BRD4 does not globally inhibit Aurora B kinase activity but rather modulates it. As there is currently no consensus on the precise role of histone H3 phosphorylation in mitosis, we cannot speculate on the differential effects of BRD4 on Aurora B phosphorylation of histone H3 isoforms. BRD4 also inhibited Aurora B phosphorylation of MCAK at S95, a prerequisite for its correct localization and activation, but BRD4 kinase activity was dispensable here. These varied effects of BRD4 on Aurora B kinase activity suggest a multifaceted modulatory mechanism.

BRD4 suppression of Aurora B activity is relieved upon JNK activation during metaphase, which phosphorylates BRD4 and releases it from chromatin (9, 10). Although JNK activity has been linked to the expression of Aurora B and the activation of its kinase through upstream mechanisms in vivo (27), the activation mechanism was unknown. Our results reveal a mechanism whereby JNK relieves the suppression of Aurora B kinase activity through its phosphorylation of BRD4, which releases it from chromatin and disrupts its interaction with Aurora B. Thus, the phosphorylation and function of Aurora B substrates such as MCAK depend on a cascade of events that begins with JNK phosphorylation of BRD4, leading to the release of BRD4 and consequent activation of Aurora B. It should be noted that Aurora B activity is not completely shut down at any stage of the cell cycle but rather peaks and dips at different stages. This is consistent with varying levels of JNK activity at different cell cycle stages and our finding that the degree of BRD4 colocalization with Aurora B varies with the different stages of the cell cycle. Notably, JNK does not directly affect Aurora B activity or its substrates in vitro. We speculate that the extent of BRD4-Aurora B interaction may vary depending on cell type, cell cycle stage, chromatin vs. non-chromatin localization and possible stress response.

Based on our results, we propose a model for BRD4 regulation of Aurora B during mitosis (Fig. 4B). Chromatin-bound BRD4 directly interacts with Aurora B, resulting in the inhibition of Aurora B kinase activity. During metaphase, JNK activity peaks, leading to its phosphorylation of BRD4 and the consequent release of BRD4 from chromatin, reducing its binding to Aurora B. The dissociation of BRD4 from Aurora B restores Aurora B kinase activity, which then phosphorylates its substrates, such as histone H3 and MCAK, to ensure orderly mitotic progression. The transiently released BRD4 returns to chromatin as a transcriptional activator that is not co-localized with Aurora B.

Limitations of the study: The precise sites of the BRD4 interaction and phosphorylation on Aurora B remain to be identified. Aurora B is part of the chromosome passenger complex (CPC), which consists of the INCEP protein that is known to tightly regulate Aurora B activity (23). Possible interplay between BRD4 and INCEP in regulating Aurora, if any, will thus need to be characterized and could increase the complexity of any regulatory network.

## Data availability

All data are contained within the manuscript. All materials and reagents will be made available upon request. The custom macros used for PLA quantification and montage generation are available upon request.

## Acknowledgements

The authors thank Drs. Hyun Park and Ananda Roy for critical reading of the manuscript and for other members of the Singer lab for ongoing discussions.

## Author contributions

DB, DC and AS conducted experiments. DB and DS designed the study and wrote the manuscript. DC helped edit the manuscript.

## Funding and additional information

This research was funded by the Intramural Research Program of the NIH, National Cancer Institute, Center for Cancer Research. This study was conducted prior to BD’s current employment at the Center for Scientific Review (CSR), NIH, and is independent of CSR.

## Experimental procedures

### Cell Lines and Culture

HCT116 were acquired from ATCC. DLD1-BRD4-IAA7 cells were a gift from Dr. Ali Shilatifard, Northwestern University, Chicago, IL. and are as described previously (17). HCT116 and DLD1-BRD4-IAA cells were grown in DMEM media with 10% FBS at 37°C and 7.5% CO2. Cell lines were tested for mycoplasma contamination. Drosophila SF9 cells were grown at 27°C in TNM-FH insect medium (BD Biosciences Pharmingen).

### Plasmid constructs

Murine 6X His-BRD4-Flag WT and ΔN, ΔC, ΔB1, ΔB2 and ΔB1B2 mutants are as described previously (Ref.). The human 3A-BRD4 mutant is as reported previously (9).

### Recombinant proteins

Flag and His tagged BRD4 and BRD4 mutants were purified from insect cells as described earlier (8). GST-Pol II CTD was purified from E.coli cells by inducing expression using 0.7mM IPTG overnight at 37°C and purified using GST Sepharose beads (Sigma) All purified proteins were concentrated using microcon size exclusion columns (Millipore), recovered in HKEG buffer (20mM Hepes, pH 7.9, 100mM KCl, 0.2mM EDTA, 20% vol/vol Glycerol) and stored frozen at - 80°C. Purified recombinant kinase active human GST-tagged Aurora B and JNK1 was purchased from SignalChem. Purified GST-MCAK kinesin motor domain protein was purchased from Cytoskeleton Inc. Purified Human histones H3 proteins were purchased from New England Biolabs. Purified recombinant human His-MYC was purchased from Raybiotech. Purified recombinant PTEFb (CDK9) was purchased from Sigma-Millipore.

### Antibodies and reagents

Antibodies used for either immunoprecipitation or immunoblotting were anti-BRD4 mouse monoclonal antibody (Sigma; AMAB90841), anti-Aurora B rabbit polyclonal antibody (Abcam; ab2254), Anti-His probe (SantaCruz biotech H-3: sc-8036), anti-Histone H3 (Cell Signaling), anti-Pan-Acetyl Histone H3 (Millipore), Anti-Histone H3 pS10 (SantaCruz biotech; sc-8656-R), Anti-Histone H3pS28 (GeneTex, GTX128953), anti-MYC and anti-MYC pThr58 (Abcam, Y69: ab32072 and ab185655), anti-CDK9 (SantaCruz biotech D-7: sc-13130), anti-CDK9 pT186 (Sigma; SAB4504223), anti-KIF2C/MCAK pS95 (LSBio; LS-C381163), anti-JNK and anti-pJNK (SantaCruz biotech D-2: sc-7345 and G-7; sc-6254) and anti-beta tubulin (Abcam, ab6046). anti-BRD4 pThr1212 was custom-made and will be shared upon request. Anisomycin (Goldbio; A-580-25) was dissolved in dimethyl sulfoxide (DMSO); JNK inhibitor D-JNKI-1 (MedChemExpress; HY-P0069) and Indole-3-acetic acid/Auxin (Sigma; I5148) was dissolved in water to make stock solutions. Stock solutions were stored at -80°C.

### Co-Immunoprecipitations and *in vitro* binding assays

To co-immunoprecipitate specific proteins from cell extracts as detailed in the figure legends, magnetic beads (Dynabeads Protein A; ThermoFisher Scientific) were coated with 5μg of the bait antibody and incubated with the cell extract for 3hr at 4°C. The beads were then washed thrice with 50mM Tris (pH 8.0), 200 mM NaCl, and 0.2% NP-40. Bound proteins were separated on SDS PAGE gels and immunoblotted with specific antibodies as mentioned in the figures. For *In vitro* binding assays, Flag-BRD4 was pre-incubated for 1hr with M2 Flag-agarose (Sigma), beads and then incubated overnight at 4°C with recombinant purified Aurora B. The beads were washed twice with 50mM Tris (pH 8.0), 150 mM NaCl, and 0.2% NP-40 and immunoblotted with antibodies against Aurora B and BRD4. All immunoblot analyses were performed using secondary antibodies from Li-Cor and the Odyssey^TM^ infrared scanner.

### In situ Proximity Ligation assays

For PLA, approximately 10^4^ HCT116 cells were grown overnight in µ-Slide Angiogenesis (I-bidi). PLA was conducted using the Duolink® In Situ PLA® Kit (Sigma) according to the manufacturer’s protocol. The primary antibodies used were as follows: anti-BRD4 mouse monoclonal antibody (Sigma; AMAB90841) (1:100 dilution), anti-Aurora B rabbit polyclonal antibody (Abcam; ab2254) (1:100 dilution). Cells were observed with Zeiss LSM880 Multi-Photon Confocal Microscope. The PLA signal quantification was done using ImageJ with a custom macro. Statistics were performed with one-way Anova using GraphPad Prism 10, where *n* = 56, ns, *P* > 0.05 and \*\*\*\**P* < 0.0001.

### Non-chromatin and chromatin-bound protein fractionation

Treated and untreated cells were subjected to the REAP protocol as described by Suzuki and colleagues (28) to collect the nuclear fraction. The nuclear pellet was resuspended in approx. 3x volume of ice-cold buffer A (10mM HEPES pH 7.9, 1.5mM MgCl2, 10mM KCl, 1mM DTT, 0.5mM PMSF and 1x protease inhibitor cocktail) with 0.5% NP-40 and 75mM NaCl, incubated on ice for 10min, followed by centrifugation at 5000g, 4°C for 5 min. The above steps were repeated twice, and the supernatants pooled to be used as the non-chromatin bound protein fraction. The remaining pellet was resuspended in approx. 4x volume high salt buffer (10mM HEPES pH7.9, 20% glycerol, 350mM NaCl, 1.5mM MgCl2, 0.4mM EDTA, 0.5% NP40, 1mM DTT, 0.5mM PMSF and 1x protease inhibitor cocktail) and rotated at 4°C for 30 min. The samples were then centrifuged at 12000g at 40C for 10 min to collect the supernatant to be used as the chromatin-bound protein fraction.

### *In vitro* Kinase assays

*In vitro* kinase assays with recombinant proteins were performed in 20 μl of 50mM Tris (pH 7.5), 5mM DTT, 5mM MnCl_2_, 5mM MgCl_2_ with 10μCi of γ^32^P ATP (6000Ci/mM) and/or 40μM ATP where indicated in the figure legend. The kinase reactions for incubated for 1 hour at 30°C, following which the proteins were resolved by SDS-PAGE and the extent of phosphorylation quantitated by a phosphorimager. When phosphorylation was determined by immunoblotting as indicated in the figures, kinase assays were performed with unlabeled ATP.

### HAT assays

HAT assays were done as described previously (20) with minor changes. Purified BRD4 (500 ng) was incubated with 1 μg of histone substrate (New England Biolabs), 0.6 mM unlabeled Acetyl CoA (Sigma) in the presence of HAT buffer (50 mM Tris pH 8.0, 1 mM DTT, 0.1 mM EDTA and 25% V/V glycerol) at 30°C for 30 min. The reaction was stopped with SDS sample buffer, and the samples were run on 15% SDS-polyacrylamide gels that were immunoblotted with the appropriate antibody as mentioned in the figure legends.

### Transient transfections

Transient transfections of the mammalian expression plasmid constructs containing BRD4 and 3A-BRD4 were done using Lipofectamine (Invitrogen) and harvested 18 hrs post transfection except were mentioned otherwise. Transfected cells were treated with 30 µg/ml Anisomycin (A) for two hours and/or 500 µM auxin for 1 hr before harvesting where indicated in the figure legends. Whole cell extracts were made from the cells and analyzed by immunoblotting using specific antibodies as indicated in the figure legends and equal amounts loaded on gels based on total protein quantities.

### Immunofluorescence analysis

HCT116 cells were plated on cover slips. The cells were fixed with 4% paraformaldeyhde, permeabilized with 0.5% Triton-X, labeled with anti-BRD4 (Sigma; AMAB90841) and anti-Aurora B (Abcam; ab2254) antibodies and stained with DAPI. Confocal images were acquired using a Zeiss LSM510 META confocal microscope. Images were taken from 27 cells in interphase, 68 cells in prophase, 38 cells in metaphase, 25 cells in anaphase, 14 cells in telophase, representative images are shown in Fig. 4.

